# Methods for High Fidelity Spectral Data Collection for Generating Ground Truth Data for Simulated Tissues

**DOI:** 10.1101/2020.02.25.964924

**Authors:** Alex T. Gong, Arjun Dulal, Matthew M. Crane, Troy E. Reihsen, Robert M. Sweet, Alexander R. Mendenhall

**Author notes:** Equal contribution (Alex T. Gong, Arjun Dulal).

## Abstract

Our research is focused on creating and simulating hyper-realistic artificial human tissue analogues. Generation and simulation of macroscopic biological material depends upon accurate ground-truth data on spectral properties of materials. Here, we developed methods for high fidelity spectral data collection using two differently colored simulated skin tissue samples and a portable spectral imaging camera. Using the standard procedure, we developed, we quantified the reproducibility of the spectral image signatures of the two synthetic skin samples under natural and artificial lighting conditions commonly found in clinical settings. We found high coefficients of determination for all measures taken under the same lighting. As expected, we found the spectral image signature of each sample was dependent on the illumination source. Our results confirm that illumination spectra data should be included with spectral image data. The high-fidelity methods for spectral image data collection we developed here should facilitate accurate collection of spectral image signature data for gross biological samples and synthetic materials collected under the same illumination source.

## INTRODUCTION

Medical error accounts for up to 250,000 deaths in hospitals annually, corresponding to an estimated total cost between $17 billion and $29 billion^1,2^. Some of this medical error can be attributed to differences in medical training. Currently, the major variables in medical training include both the instructor and the biological sample materials available. In order to reduce training variability due to differences in instructors, there has been a gradual shift in medical training from the traditional apprenticeship approach^3^ to simulation based medical education and training^4^. Simulation-based training utilizing both low- and high-fidelity physical^5–8^ and virtual (AR/VR)^9–18^ simulation, has proven useful in improving both technical and non-technical skills. Studies done in the field of advanced cardiac life support^19,20^, cardiac auscultation^21,22^, laparoscopic surgery^23–25^, central venous^26,27^, hemodialysis catheter insertion^28^ and thoracentesis^27,29^ show that the implementation of simulation-based medical education with deliberate practice is superior to the apprenticeship approach. The two main goals contributing to this shift are: 1) improving the effectiveness and consistency of medical training, and 2) reducing the reliance on gross specimen (animal and/or human) availability for training.

These simulators require accurate ground truth data on the medical procedures they are simulating, including data on the physical properties of the biological materials being simulated. Accurate characterization of the constitutive properties of human soft tissues remains a gap that has prevented the rapid expansion of synthetic human tissue modeling^30^. Additionally, physically simulated gross specimens provide trainees a consistent training tool, ensuring that all trainees are exposed to the same samples, some of which may be exceedingly rare to encounter, if not simulated. Therefore, we seek to develop both a spectrum of synthetic training materials, and an array of AR/VR training programs. One way to achieve both of these goals is to develop synthetic materials for simulation and implement them in appropriate curricula for medical trainees. Thus, our goals are to generate a comprehensive array of accurately simulated gross anatomical analogues and corresponding AR/VR digital models. We believe that exposing the students to the same training and the same spectrum of medical samples should reduce the disparities in medical training. We also believe that we can democratize the expertise of the most renowned practitioners by digitizing every aspect of their training programs possible into AR/VR environments. Hence, by generating synthetic training materials, we free ourselves from the uncertainty of specimen availability; and by digitizing the training programs we increase access and provide more students higher quality training experiences. The overall goal is to reduce the disparities in medical training and come out with a more standardized set of trainees, in terms of medical capacitates learned from a shared training experience.

In order to simulate biological tissues in physically or digitally, we need to collect accurate data on the physical properties of those tissues. Within the last century, the mechanical properties of human tissue have undergone study^31–52^. And, while much work has been done attempting to ascertain the spectral properties of tissues in relation to pathological conditions, much remains to be learned as our technological capacities continue to improve^53^. Optical properties are important to accurately simulate biological material colors, and heterogeneity in color, in order to provide the necessary cues for healthcare providers to recognize, diagnose and treat healthcare conditions. Prior studies on the optical properties of tissues are focused primarily on optical properties such as scattering, absorption, fluorescence, and refractive index.

Currently, there is no standard for quantifying the emission spectra of real or simulated biological materials. There is also no standard for quantifying the effect of illumination on the sample spectrum. The only medical illumination standard is the IEC 60601^54^, which refers to the safety requirements for surgical luminaries and luminaries for diagnosis. There is also the CIE Δ E 2000, which is a standard for comparing color differences in a 2D color space consisting of three different axes (black to white, green to red, and blue to yellow), as in Lee et al. 2005^55^. But, this CIE Δ E 2000 and its predecessors do not directly use spectral data; they compare converted spectra in color space, derived from the original CIE Δ E in 1976. With spectral data collection taken on the same high-fidelity camera (see Figs. 1&2), this valuable standard is not necessary for direct comparison of spectral waveform data. Every color we see is comprised of some waveform spanning the visible and near infrared radiation spectrum, collected in the same 7nm bins. Thus, one more direct approach for comparing different visible light spectral image signatures is direct comparison of the spectral waveform using correlations between the spectral bin values from two different samples.

**Figure 1:**
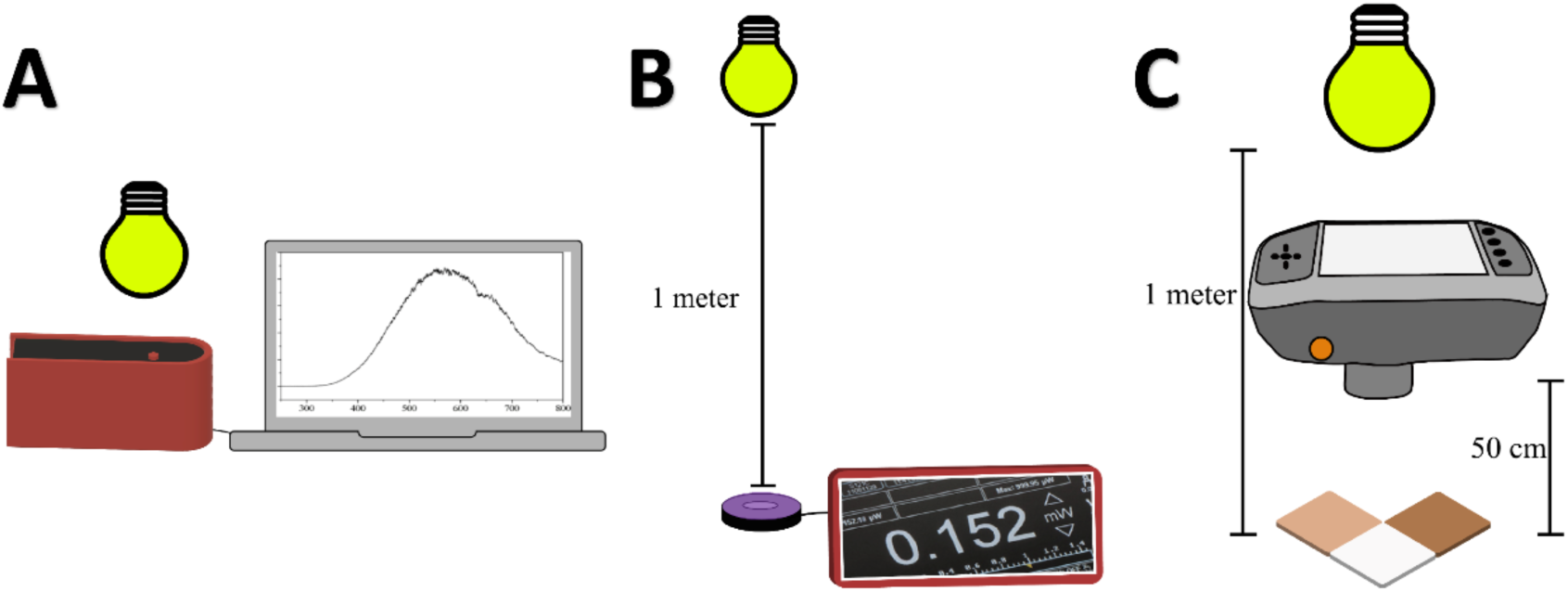
Data collection scheme. The experimental setup for collecting a) the emission spectrum, b) the illumination power, and c) the scattered spectrum.

**Figure 2:**
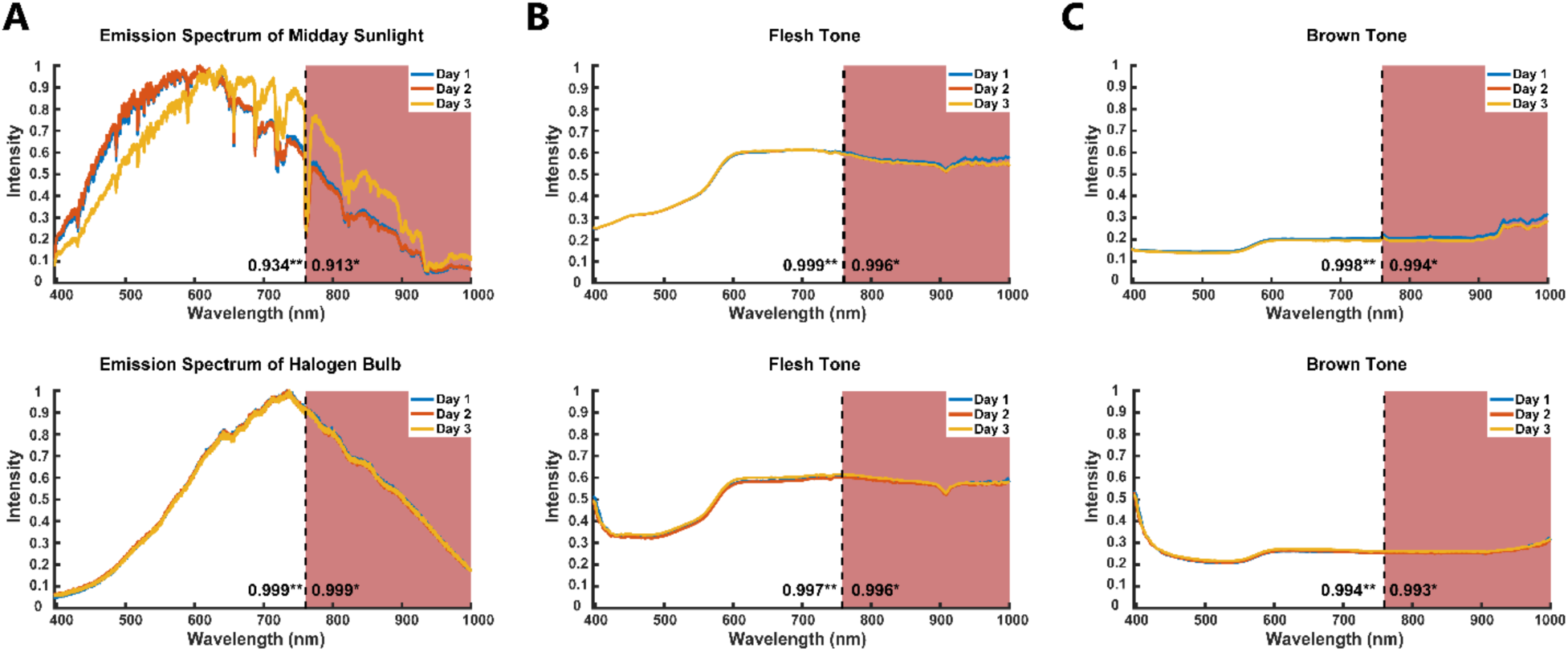
Reproducibility studies across three days of a) the emission spectrum of midday sunlight and halogen light, b-c) emission spectra of two different synthetic skin samples under midday and halogen light. (r*) and (r**) represents the average R^2^ between day 1 to day 2, day 2 to day 3, and day 1 to day 3 for the full-spectrum (400- 1000 nm) and the isolated visible spectrum (400-760 nm), respectively. The R^2^ of the illumination sources and samples indicate high reproducibility in our test setup.

We needed to develop a methodology to reproducibly collect the most accurately possible spectral data for real and synthetic tissues. A reasonably accurate way to collect data on the color of an object is to measure the spectrum of the scattered light using devices that generate spectral images^56^. Until 2017, these devices were not completely self-contained and portable^57^, needing to be attached to other equipment like a computer. Now, we are able to acquire spectral data using a completely portable, self-contained spectral camera. We developed procedures for acquiring spectral data on synthetic or real biological specimens in different medical environments. We surveyed a variety of medical lighting environments and found significant differences between the illumination spectra, both relative to one another and to the sun. We found that the scattered spectra of synthetic skin samples were extremely reproducible in the same illumination environment, with trial-to-trial variation attributable mostly to differences in near-infrared radiation (760-1000nm)^58^. We also found that the different illumination sources in different medical environments significantly affected the scattered spectra of our skin samples. In this study, our goal is to develop means to collect reproducible, accurate ground truth data on the scattered visible light spectra of physical materials for the purposes of medical training and medical simulation.

## RESULTS

### A standardized measurement scheme results in reproducible spectral data

In order to quantify data in a reproducible fashion, we devised standard protocols for collecting data on illumination spectra, illumination radiant flux, and material spectra. To collect the emission spectrum of each light source, we placed the emission spectrometer (Thor Labs, Newton, New Jersey) close enough to the light source to clearly detect the features of each emission spectrum, but not saturate the detector – usually within a few centimeters of the light source. To quantify the radiant flux from an illumination source, we placed a power meter (Thor Labs, Newton, New Jersey) with an emission filter (Leica Microsystems, Wetzlar, Germany) one meter from each light source and measured the radiant flux. After measuring the radiant flux with the emission filter, we then calculated the total radiant flux of the entire emission spectrum from our sample, using the power sampled at discreet bandwidth and MatLab’s TrapZ function (see Methods). To collect the scattered light emission spectra of synthetic skin samples, we placed each skin sample one meter directly under the light source, and then we mounted a spectral camera at a 5-degree angle facing the sample, and acquired an image, which took between a few seconds and a minute, depending on the intensity of illumination. Figure 1 shows our overall data collection scheme for illumination radiant flux and spectrum, and sample spectrum.

We evaluated the reproducibility of our methodology by quantifying the correlations between spectra measured on different days as shown in Figure 2. Supplementary Figure 1a and Supplementary Table 1 shows the reproducibility of all light sources quantified. We found that the 5000K 99 CRI LED and the halogen bulb illumination sources were most consistent (R^2^=1.000, 3nm spectral resolution) and the sunlight Illumination was least consistent (R^2^=0.9132) across the spectrum. Here we collected illumination and sample scattered spectral data as shown in Figure 1. Reproducibility scattered spectra of two synthetic skin samples under midday sun and halogen lighting conditions are shown in Figure 2b,c. Supplemental Figure 1b,c shows the reproducibility of each of our two synthetic skin samples under all the lighting conditions. Supplementary Table 2 and 3 lists the correlations between each sample across three days of measurements under each of eight illumination conditions, with and without infrared radiation considered. Overall, the 5000K 99 CRI LED illumination provided the most consistent measurements of synthetic skin spectrum, across the full-spectrum, for both samples (R^2^=1.000) compared to the fluorescent illumination which provided the least R^2^ values of 0.8358 and 0.9332 for Samples 1 and 2, respectively. The inter-measurement variation in scattered sample spectra is driven by differences in near-infrared radiation, shown by increased correlations when removing infrared spectral data from consideration, in Supplementary Tables 2 and 3.

### Common medical illumination spectra are distinct from both the sun and each other

We collected the emission spectrum of illumination sources and the scattered spectrum of flesh tone and brown synthetic skin samples. For illumination, we collected the emission spectrum and radiant flux for midday sunlight, halogen bulbs (commercial and operating room (OR) approved), fluorescent lights, 3000K LEDs, 4000K LEDs, 5000K LEDs, and 5000K 99 CRI LEDs. To compare the spectra, we normalized each curve to peak visible light intensity, as shown in Figure 3. The radiant flux emitted by the sun exceeded the detection limit of the power meter used in this study. Kopp et. al.^59^, measured total solar irradiance during the 2008 solar minimum period to be 1360.8 +/- 0.5 W/m^2^ ^59^ or 96.46 mW as calculated based on the diameter of our detector.

**Figure 3:**
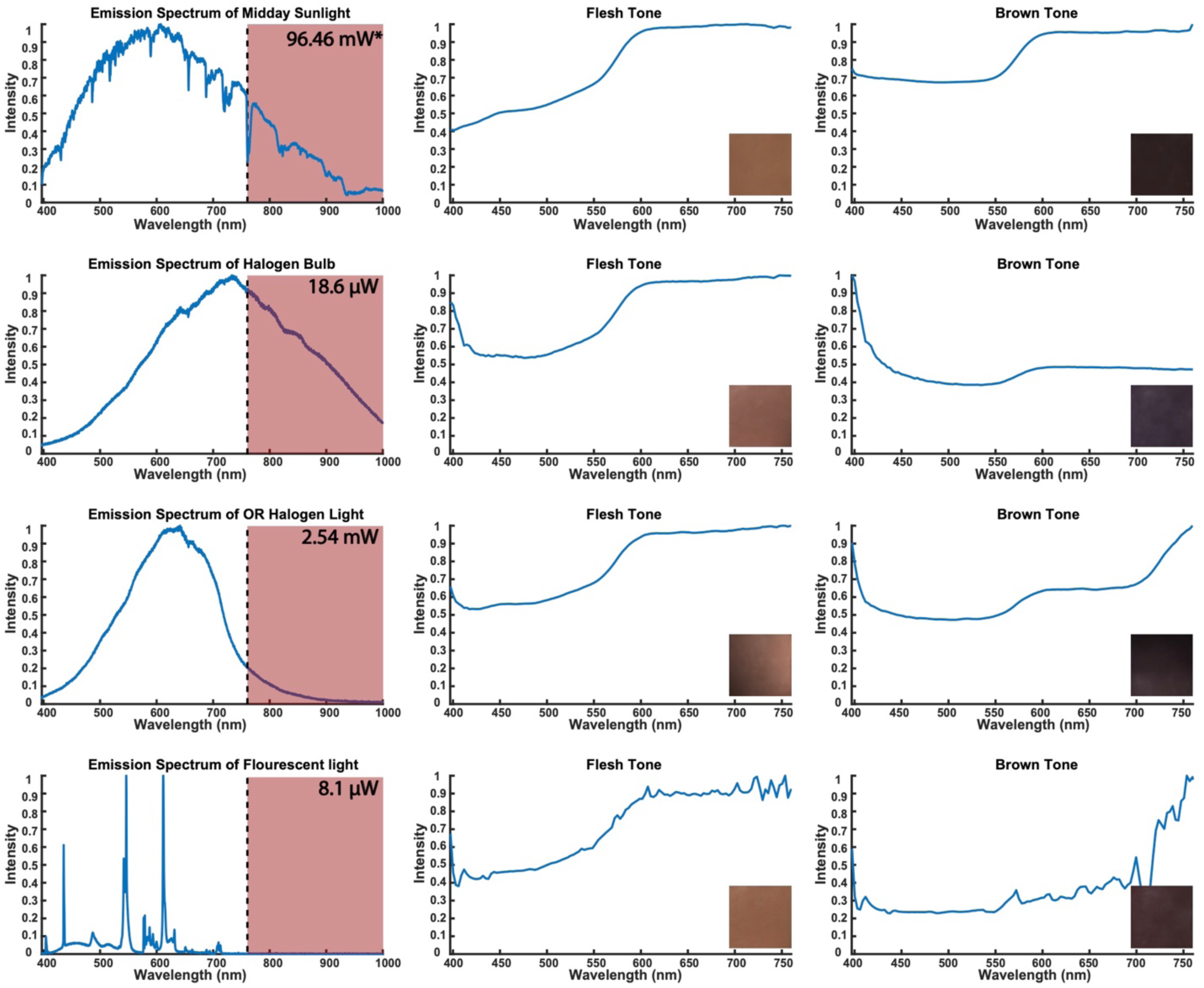
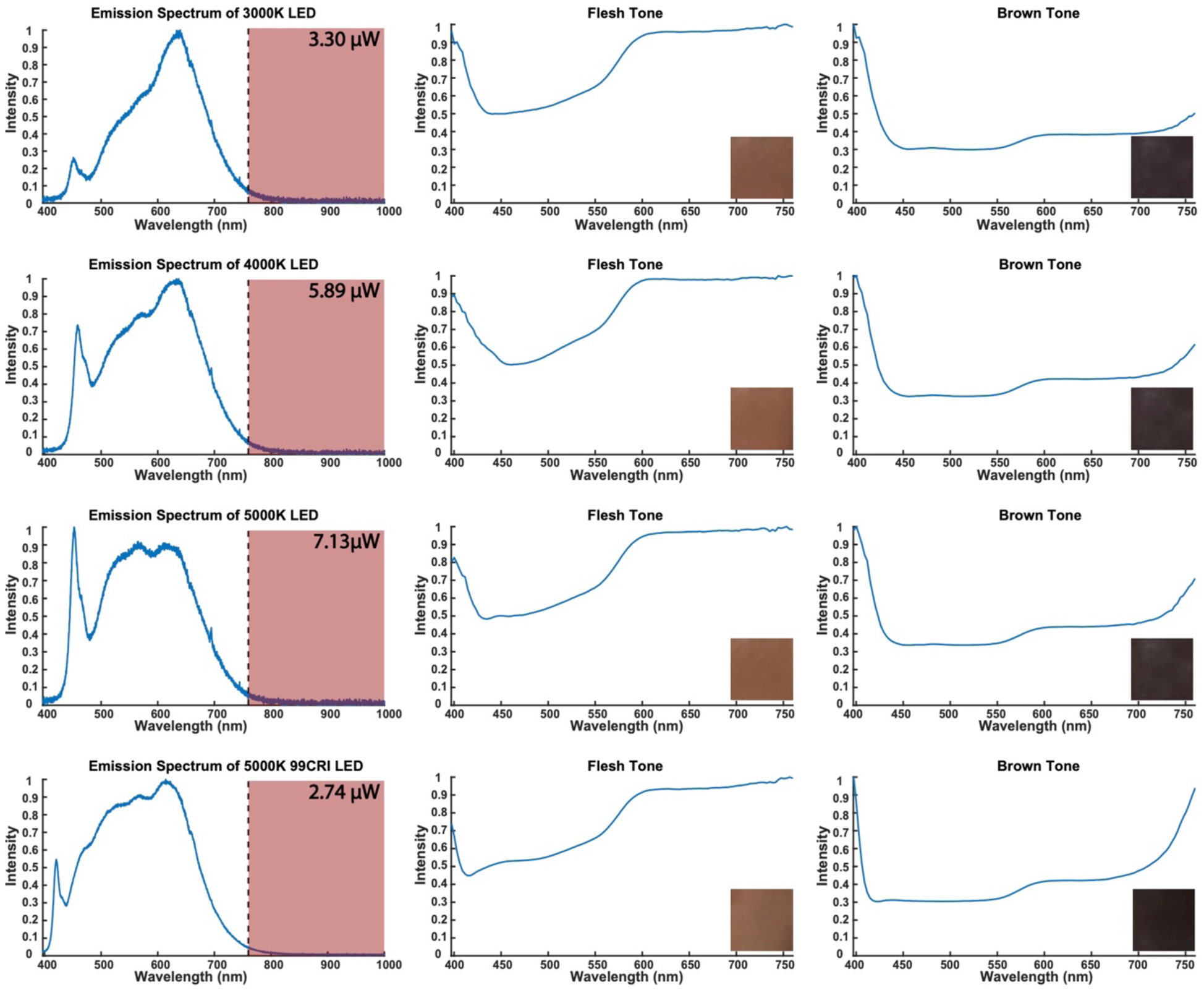
Raw emission spectrums and corresponding power, sample scattered spectrums and spectral image were quantified. The shaded region indicates the infrared regime. The sample spectrums were normalized within the visible spectrum (400 – 760 nm).

Figure 4 and Supplementary Table 4 shows the correlation between the emission spectra for all light sources and the sun. We note that our instrumentation accurately detected previously reported dips, attributed to chemicals in the atmosphere, in the generally parabolic sun spectrum^60^. We found the 4000K LED lights produced an emission spectrum most similar to the sun (R^2^ = 0.8271) when analyzing within just the visible spectrum; whereas, when expanding to the full-spectrum, the operating room halogen lights dominate (R^2^ = 0.8030), followed closely by the 4000K LED lights (R^2^ = 0.7933).

**Figure 4:**
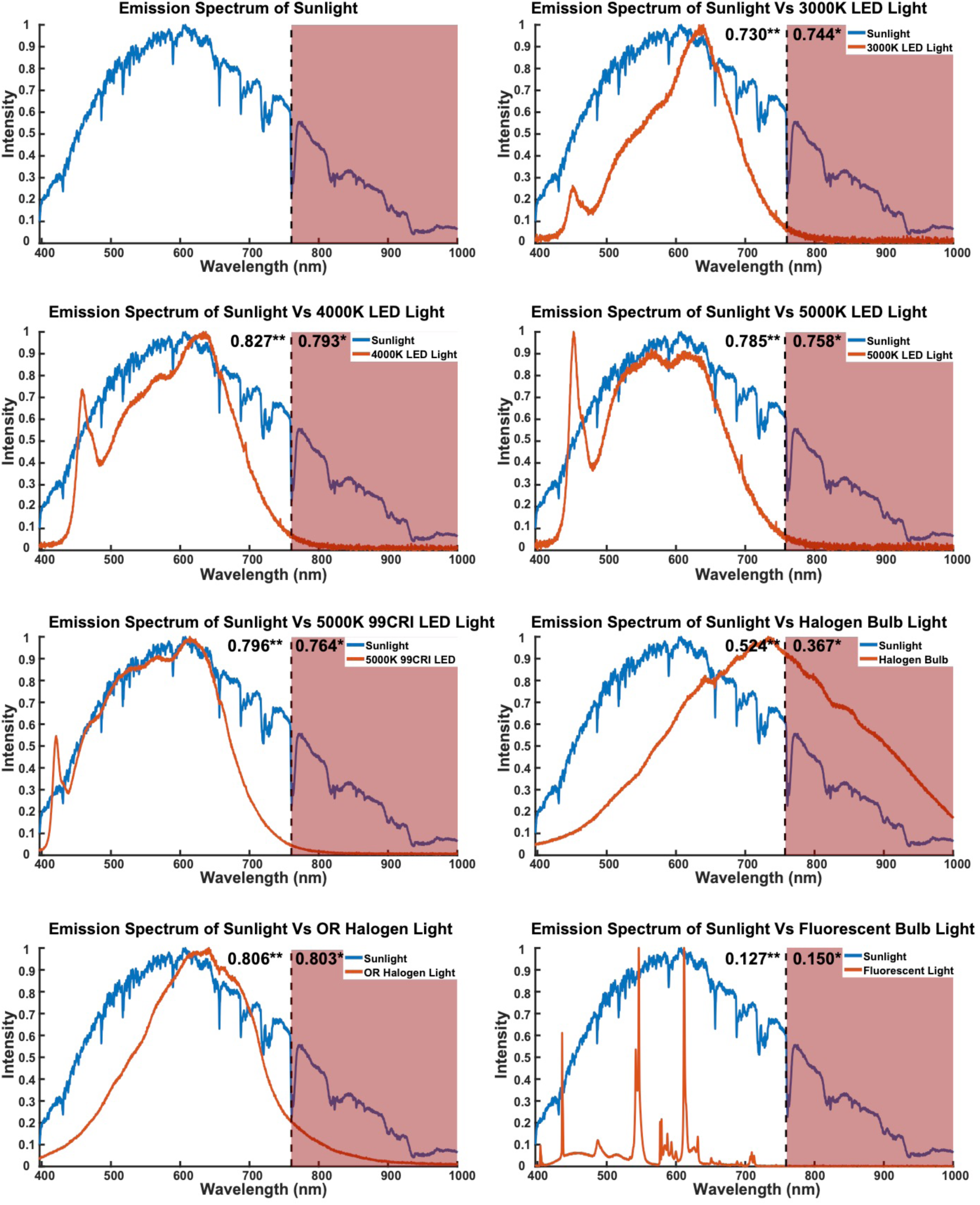
The emission spectrums of light sources were compared to the sun to assess the full-spectrum capabilities of artificial lighting. The halogen emission spectrum replicates the full-spectrum nature of sunlight (R^2^=0.8030) the most, whereas fluorescent lighting (R^2^=0.1504) is the least similar. The variation in emission spectrums leads to quantitative and qualitative differences in the perception of color.

### As expected, but important to demonstrate, common medical illumination sources have distinct effects on spectral image signatures on differently colored synthetic skin

Figure 5 shows the correlation between the scattered spectra of the synthetic skin samples under various lighting conditions compared to spectra scattered by the sun. Supplementary Table 5 shows the correlations between the scattered spectrum of each sample under each corresponding lighting condition. Synthetic flesh tone skin looked spectrally most similar to the sun when illuminated by just the visible spectrum, parabolic OR halogen lights (R^2^ = 0.9794), which also had the most watts of power cast upon the sample (2.54 mW). When expanded to the full-spectrum, flesh tone was most correlated under the commercially available halogen bulb. Synthetic brown skin tone looked most spectrally similar to the sun when restricted to the visible regime under the OR halogen lights (R^2^ = 0.5702). When expanded to include the full-spectrum, the OR halogen lights (R^2^ = 0.7813) elicited the highest correlated scattered spectrum to the sun. The synthetic flesh tone samples consistently exhibited a blue shift when compared to the sun; whereas, the synthetic brown tone samples resulted in both a blue and red shifted.

**Figure 5:**
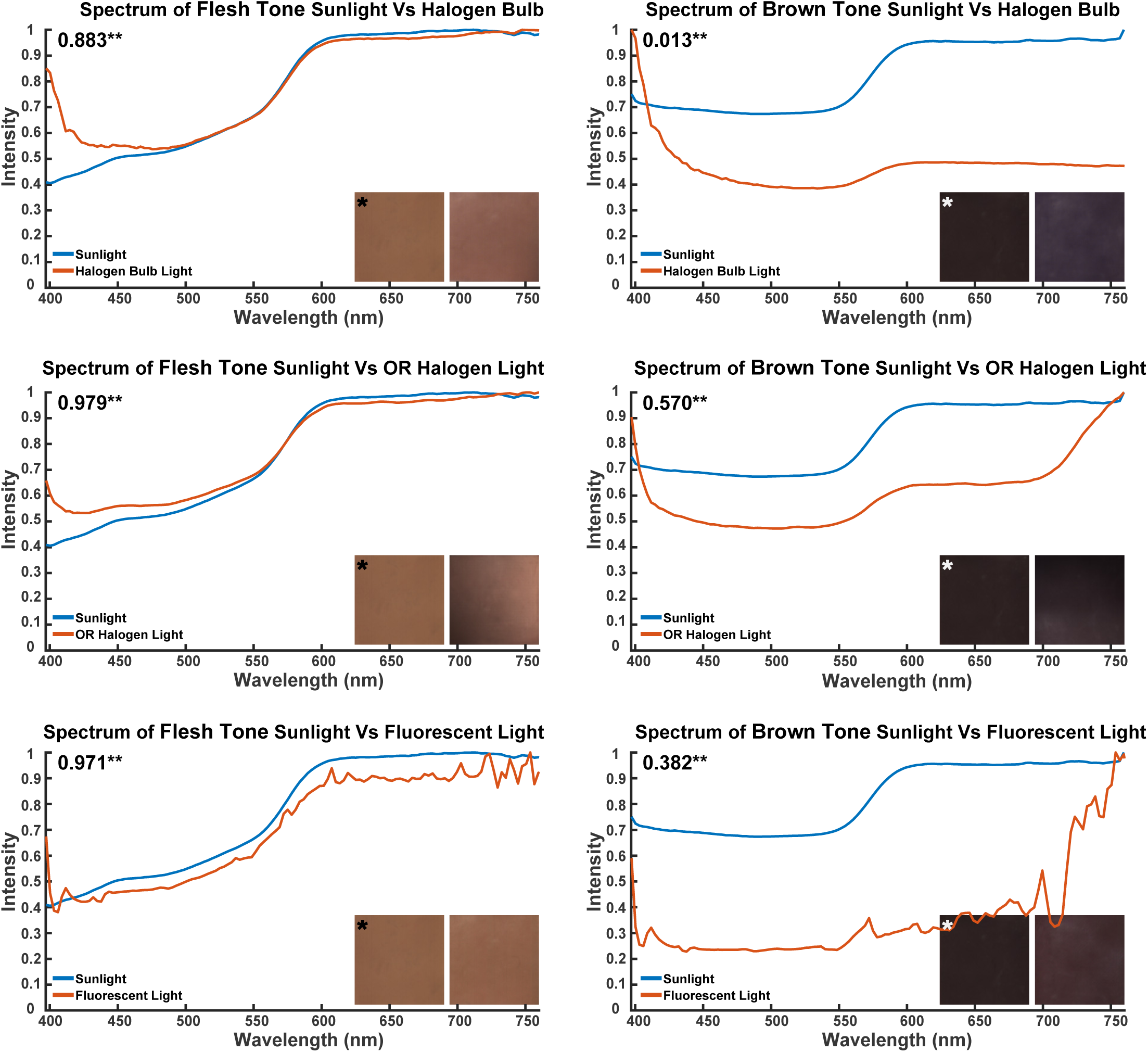
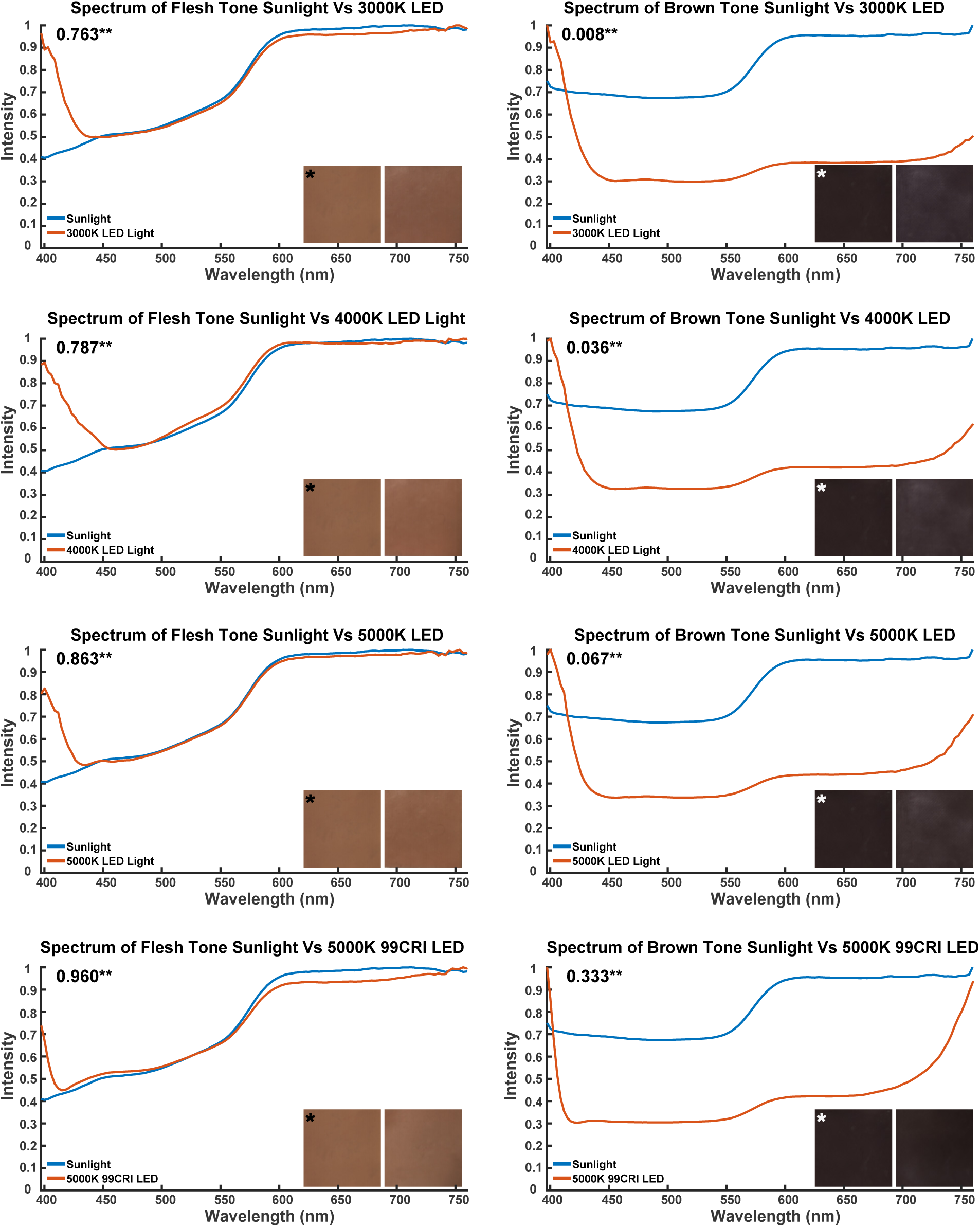
Scattered spectra of two synthetic skins were compared under eight artificial light sources to the sun. The spectral image of the sun and test sample show visible color differences. The R^2^ of each sample as compared to each lighting condition is indicated by the double asterisk. As expected, the highest correlation between the sun and artificial lighting was the full-spectrum halogen bulb. Interestingly, all artificial lighting displayed poor correlation to the sun on the darker sample, with the highest correlation coming from the halogen bulb.

## Discussion

### Illumination in medical practice

The IEC 60601^54^ standard for surgical luminaries allows physicians to operate in 3000-5000k lighting conditions. Given the difference in synthetic skin sample scattered spectra for LEDs between 3000 and 5000k in color temperature (Figure 3), this will have significant effects on the emission spectra of patients. If the scattered spectra of patient biological materials are different, the clinicians’ ability to perceive physiological conditions may be different. Thus, we strongly suggest considering the types of observations necessary for a particular medical procedure, including the colors necessary to be perceived, when selecting the color temperature of light used in medical practice.

### Standards for acquiring biological material spectrum data

While there are currently no standards for acquisition of tissue color data, we suggest that researchers start data collection using a full spectrum light source. In the past year, full spectrum LEDs that mimic the solar emission spectrum (Gamma Technologies, San Diego) have become commercially available. Our research indicates that the emission spectrum of the illumination source should be presented with the scattered spectrum of the sample material. We suggest the radiant flux on the sample be recorded in order to understand how a measurement may be influenced by the amount of illumination in which it was bathed. Thus, we should include the illumination source spectrum in order to understand any spectral discrepancies that may arise under other lighting conditions. Furthermore, we note that the variation in spectral data collection was often influenced by radiation in the near-infrared, and that the near-infrared spectrum should be reported and considered.

### Perceived and real differences in color in medical practice and training

Perception of color is important for both practice and training. In practice, one needs to be able to discriminate between shades of red (flushing), yellow (jaundice), and blue (hypoxia/hypothermia). Conversely, when training to recognize these changes in patient state, it is critical that the trainees be exposed to normal and abnormal tissues, across the spectrum of human skin tones. It is important to note the aforementioned color recognition-based diagnoses are on the surface of the body and the potential for the effects of metamerism. Once inside and performing laparoscopic or open-cavity surgery, there are a myriad of other color-indicated changes in physiological state to observe. We plan to digitally codify these changes and ensure that they are accurately represented to a much broader group of trainees than would be possible with simple gross specimens or trainee-observed medical procedures.

In this paper we showed both qualitative and quantitative markers indicating differences in the perception of color under eight different lighting conditions. Artificial lighting conditions have been shown to introduce optical differences within samples when compared to sunlight. For example, the LED lighting on brown tone, created a blue shifted spectrum. In addition, the initial color of the sample itself can affect the degree of color change between lighting conditions. The flesh tone, on average, exhibited a significantly higher degree of linear correlation in terms of its scattered spectrum when compared to the sun; whereas the scattered spectra of the brown tone was minimally or not correlated between different illumination spectra. This suggests there are intrinsic differences in perception disparities solely based on the fact that different illuminations affect the spectral shifts of different colors in significantly different ways (e.g. See Figure 5a, sun vs. fluorescent bulb for flesh-tone vs. brown).

### Simulation of accurately colored synthetic materials

In order to simulate healthy and diseased biological materials for medical training appropriately, accurate color data is a necessity. Such data requires the need for using full spectrum lighting to record all the spectral features of a tissue – more than would be possible with spectrally incomplete lighting. For example, shining a blue light on a tissue sample and then recording the emission spectra would illicit a bluer spectrum, because the only photons available were blue. This would yield incomplete color data, resulting in a poor optical representation of tissue. We will be able to develop high-fidelity physical and virtual training models by collecting data under consistent full spectrum lighting conditions. By simulating tissues that exhibit rare anatomic pathologies, medical students and clinicians will be able to train and maintain necessary skills

### Simulation of accurate color in AR/VR environments

A significant task for the digitization of spectral data will be the conversion of spectral data into standard Red/Green/Blue (sRGB) values for simulation of biological tissues in digital AR/VR environments. Thus, we plan to perform research to quantify the difference between sRGB representations of biological materials and true spectra. Given how poorly sRGB LEDs mimic the parabolic spectrum of the sun or halogen lights, a likely outcome of this research may be that certain aspects of color are not reproducible with three color LED arrays. As we noted above, there are lighting companies (e.g., Gamma Scientific, San Diego) that make LED arrays that mimic the full spectrum of the sun using many different colors of LED. It may be possible to represent truer colors by using arrays of several multi-color LEDs per unit pixel in monitors. If this becomes possible, then spectral data may be more accurately represented by monitors and in AR/VR environments than would be possible with only three colors of LED.

### Implications for accurately colored materials and simulators in medical training and practice

Provided with the ability to present many different students the same materials, we believe that medical instructors will be able to more consistently produce well-trained medical practitioners. We hope that by providing a spectrum of standardized, hyper-realistic synthetic tissue samples, in conjunction with valid simulation curriculum, we can improve the quality of medical trainees by reducing variation in training and providing the visual cues necessary that drive diagnosis and decision making. Thus, it is our goal to reduce heterogeneity in medical training in order to improve the quality and consistency of modern medical practice by reducing medical error attributable to differences in training.

## Materials and Methods

### Acquisition of Illumination Spectra and Radiant Flux

We used a spectrometer (CCS200, Thorlabs, Inc., USA) in conjunction with a cosine corrector (CCSB1, Thorlabs, Inc., USA) to quantify the emission spectrum of multiple light sources. The following light sources were used in this study: 3000K 97CRI LED bulb (BC Series, Yuji International Co., Ltd, China) 4000K 97 CRI LED Bulb (BC Series, Yuji International Co., Ltd, China), 5000K 97CRI LED Bulb (BC Series, Yuji International Co., Ltd, China), 5000K 99CRI LED strip (Absolute Series, Waveform Lighting, USA), Fluorescent bulb (ED25-V, Ottlite Technologies Inc., USA), Halogen bulb (Sunbeam, USA), Halogen based surgical overhead light (PRX 4401SAX, ALM, USA) and natural mid-day sunlight. We placed the spectrometer 1m perpendicular to the light sources and captured the emission spectrum of each light source (Fig. 1b). Once the spectrum was recorded, we replaced the spectrometer with a power meter (PM200, Thorlabs, Inc., USA) with attached emission filter (ET CFP – M205FA/M165FC, Leica, USA). Using the power meter, we recorded the radiant flux of all the light sources between 440-520 nm. Finally, we use calculated the total radiant flux across the emission spectrum of each light source using the trap-z function in MATLAB.

### Synthetic Skin Swatch Synthesis

Two 100mm x 100mm x 3mm PDMS swatches were made using Platsil-Gel 25 (PolyTek Development Corp., USA) colored with 1 mL of Brown and Flesh-Tone pigments (Smooth-on, Inc., USA). To prevent contamination and particle build-up, we stored the samples in individual containers at room temperature until imaging occurred.

### Sample Scattered Spectra

We placed the skin swatch samples and white reference under each lighting condition and used the SPECIM hyperspectral camera (Specim, Spectral Imaging Ltd., Finland) to photograph the synthetic skin samples. To maintain consistency, we placed the samples perpendicular to the light sources at a distance of 1 m, except for the sunlight condition where they were positioned approximately 1.496E+11m away. We placed the hyperspectral camera at a 5-degree angle between the light source and the sample at a distance of 50 cm in such a way that the camera did not cast a shadow over the samples (Figure 1a). The camera provides a spectrum for each individual pixel in the captured image.

### Image Data Acquisition

We analyzed the scattered spectrums of the samples using a software framework for hyperspectral data exploration and processing in MATLAB^61^. We modified the script to average the pixel-based spectra from within the user selected ROI and output a normalized scattered spectrum for each of the samples. The final spectrum of each of the samples ranged from 397 nm to 1000 nm wavelength.

### Correlation Analyses for reproducibility and similarity of illumination and spectral image signature

In order to calculate the reproducibility and similarity within a light source emission spectra and sample scattered spectra, we collected the data for three days. We collected the data for each spectrum in three independent experiments executed on three different days. After collecting the data, we performed linear regression analysis in MATLAB to determine the goodness of fit between the day-to-day source emission spectra as well as corresponding sample scattered spectrum. The same regression analysis was used to determine the similarity between artificial and sunlight emission spectrum and their scattered sample spectra.

## Author Contributions

RS, TR, AG, AD and AM designed the study. AD and AG collected the data. AG, AD, MC and AM analyzed the data. AD generated plots of data. RS, TR, AG, AD, MC and AM wrote and revised the manuscript.

---

**Supplementary Figure 1:**
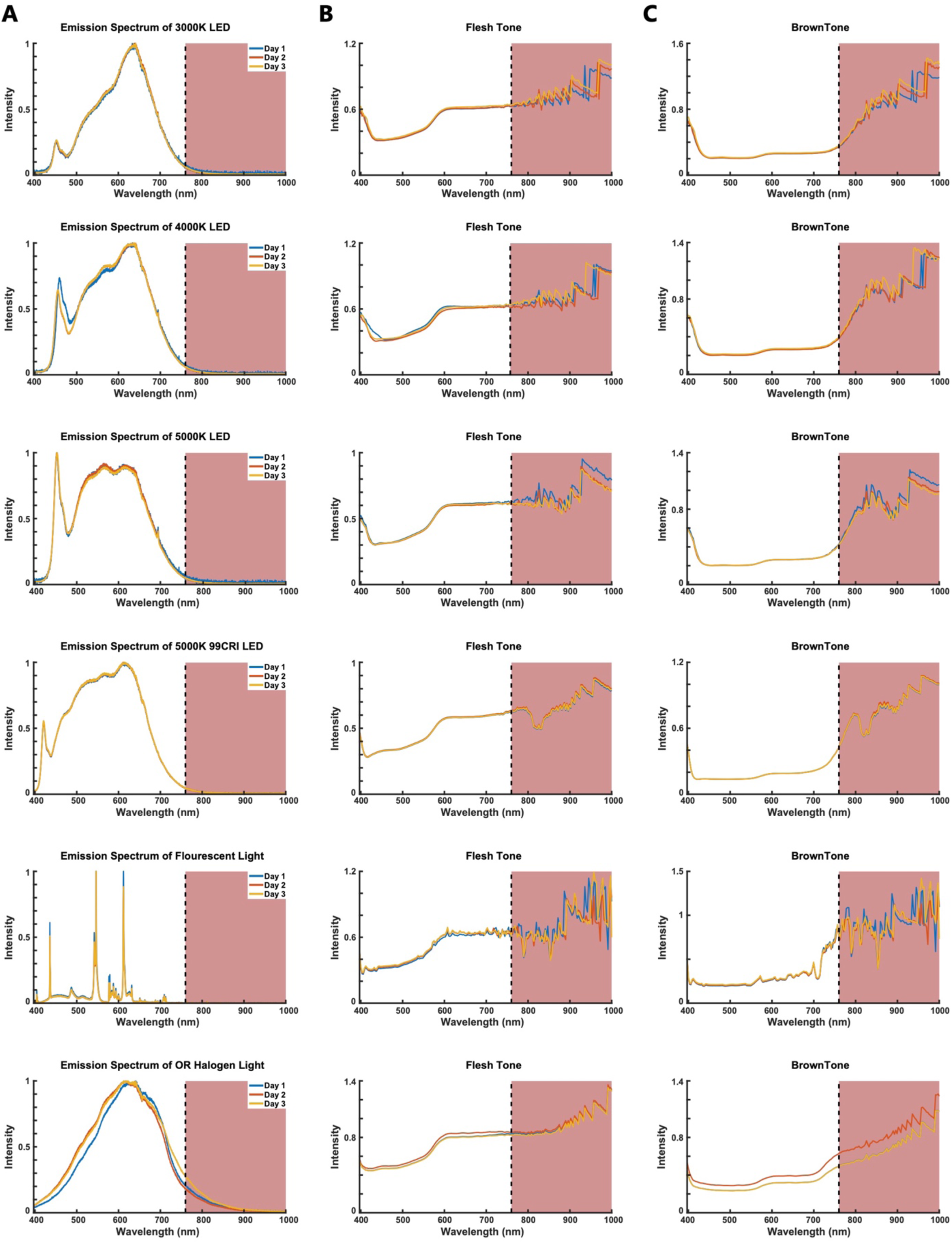
Reproducibility studies across three days of a) the illumination spectrum of light sources, b-c) scattered spectra of two different synthetic skin samples under the respective light sources. r* and r** represents the average R^2^ between day 1 to day 2, day 2 to day 3, and day 1 to day 3 for the full-spectrum (400-1000 nm) and the isolated visible spectrum (400-760 nm), respectively. The R^2^ of the illumination sources and samples indicate high reproducibility in our test setup.

**Supplementary Table 1.** Reproducibility/Similarity of the emission spectrum of light sources, within the full (*) and visible (**) range of wavelength, collected between 3 different days.

| Source | Day 1-2 | Day 1-3 | Day 2-3 | Average |  |
| --- | --- | --- | --- | --- | --- |
| **3000K LED** | 0.9991 | 0.9992 | 0.9998 | 0.9994 | * |
|  | 0.9992 | 0.9992 | 0.9997 | 0.9994 | ** |
| **4000K LED** | 0.9903 | 0.9912 | 0.9998 | 0.9938 | * |
|  | 0.9880 | 0.9891 | 0.9998 | 0.9923 | ** |
| **5000K LED** | 0.9992 | 0.9989 | 0.9997 | 0.9993 | * |
|  | 0.9992 | 0.9988 | 0.9997 | 0.9992 | ** |
| **5000K 99CRI LED** | 0.9999 | 0.9999 | 0.9999 | 0.9999 | * |
|  | 0.9999 | 0.9998 | 0.9998 | 0.9998 | ** |
| **Fluorescent** | 0.9921 | 0.9932 | 0.9994 | 0.9949 | * |
|  | 0.9919 | 0.9930 | 0.9994 | 0.9948 | ** |
| **Halogen Bulb** | 0.9994 | 0.9996 | 0.9995 | 0.9995 | * |
|  | 0.9997 | 0.9997 | 0.9997 | 0.9997 | ** |
| **OR Halogen** | 0.9813 | 0.9893 | 0.9911 | 0.9872 | * |
|  | 0.9784 | 0.9882 | 0.9914 | 0.9860 | ** |
| **Sunlight** | 0.9986 | 0.8804 | 0.8606 | 0.9132 | * |
|  | 0.9989 | 0.9096 | 0.8949 | 0.9345 | ** |

**Supplementary Table 2.** Reproducibility/Similarity of the scattered spectrum of flesh tone under the light sources, within the full (*) and visible (**) range of wavelength, collected between 3 different days.

| Source | Day 1-2 | Day 1-3 | Day 2-3 | Average |  |
| --- | --- | --- | --- | --- | --- |
| 3000K LED | 0.8945 | 0.8925 | 0.9464 | 0.9111 | * |
|  | 0.9972 | 0.9962 | 0.9954 | 0.9963 | ** |
| 4000K LED | 0.9382 | 0.8903 | 0.8663 | 0.8983 | * |
|  | 0.9843 | 0.9847 | 0.9981 | 0.9890 | ** |
| 5000K LED | 0.9610 | 0.9587 | 0.9886 | 0.9694 | * |
|  | 0.9986 | 0.9986 | 0.9995 | 0.9989 | ** |
| 5000K 99 CRI LED | 0.9996 | 0.9999 | 0.9999 | 0.9998 | * |
|  | 0.9999 | 0.9999 | 1.0000 | 0.9999 | ** |
| Fluorescent | 0.8224 | 0.8128 | 0.8723 | 0.8358 | * |
|  | 0.9786 | 0.9888 | 0.9981 | 0.9885 | ** |
| Halogen Bulb | 0.9967 | 0.9945 | 0.9971 | 0.9961 | * |
|  | 0.9965 | 0.9958 | 0.9990 | 0.9971 | ** |
| OR Halogen | 0.9986 | 0.9998 | 0.9988 | 0.9991 | * |
|  | 0.9991 | 0.9999 | 0.9996 | 0.9995 | ** |
| Sunlight | 0.9947 | 0.9947 | 1.0000 | 0.9965 | * |
|  | 0.9997 | 0.9996 | 1.0000 | 0.9998 | ** |

**Supplementary Table 3.** Reproducibility/Similarity of the scattered spectrum of brown tone under the light sources, within the full (*) and visible (**) range of wavelength, collected between 3 different days.

| Source | Day 1-2 | Day 1-3 | Day 2-3 | Average |  |
| --- | --- | --- | --- | --- | --- |
| 3000K LED | 0.9573 | 0.9569 | 0.9793 | 0.9645 | * |
|  | 0.9951 | 0.9918 | 0.9945 | 0.9938 | ** |
| 4000K LED | 0.9788 | 0.9598 | 0.9469 | 0.9618 | * |
|  | 0.9956 | 0.9975 | 0.9952 | 0.9961 | ** |
| 5000K LED | 0.9887 | 0.9893 | 0.9968 | 0.9916 | * |
|  | 0.9960 | 0.9978 | 0.9987 | 0.9975 | ** |
| 5000K 99 CRI LED | 0.9998 | 1.0000 | 0.9999 | 0.9999 | * |
|  | 1.0000 | 1.0000 | 1.0000 | 1.0000 | ** |
| Fluorescent | 0.9297 | 0.9194 | 0.9504 | 0.9332 | * |
|  | 0.9903 | 0.9914 | 0.9991 | 0.9936 | ** |
| Halogen Bulb | 0.9910 | 0.9940 | 0.9945 | 0.9932 | * |
|  | 0.9918 | 0.9956 | 0.9951 | 0.9942 | ** |
| OR Halogen | 0.9969 | 1.0000 | 0.9967 | 0.9979 | * |
|  | 0.9968 | 1.0000 | 0.9967 | 0.9978 | ** |
| Sunlight | 0.9915 | 0.9926 | 0.9999 | 0.9947 | * |
|  | 0.9979 | 0.9976 | 1.0000 | 0.9985 | ** |

**Supplementary Table 4.** Reproducibility/Similarity of the emission spectrum artificial light sources, within the full and visible range of wavelength, compared to the natural sunlight.

| Sources | Full Spectrum | Visible Spectrum |
| --- | --- | --- |
| **Sun-3000K LED** | 0.7304 | 0.7442 |
| **Sun-4000K LED** | 0.7933 | 0.8271 |
| **Sun-5000K LED** | 0.7586 | 0.7856 |
| **Sun-5000K 99CRI LED** | 0.7644 | 0.7968 |
| **Sun-Fluorescent** | 0.1504 | 0.1275 |
| **Sun-Halogen Bulb** | 0.3672 | 0.5244 |
| **Sun-OR Halogen** | 0.8030 | 0.8068 |

**Supplementary Table 5.** Reproducibility/Similarity comparison of the scattered sample spectrum produced by artificial light sources, within the full (*) and visible (**) range of wavelength, compared to the sample spectrum produced by natural sunlight.

| Source | Flesh Tone | Brown Tone |  |
| --- | --- | --- | --- |
| Sun-3000K LED | 0.5218 | 0.6140 | * |
|  | 0.7629 | 0.0088 | ** |
| Sun-4000K LED | 0.5352 | 0.6145 | * |
|  | 0.7871 | 0.0362 | ** |
| Sun-5000K LED | 0.6175 | 0.6136 | * |
|  | 0.8630 | 0.0672 | ** |
| Sun-5000K 99CRI LED | 0.6630 | 0.6723 | * |
|  | 0.9605 | 0.3338 | ** |
| Sun-Fluorescent | 0.5426 | 0.6300 | * |
|  | 0.9707 | 0.3823 | ** |
| Sun-Halogen Bulb | 0.8772 | 0.0600 | * |
|  | 0.8830 | 0.0133 | ** |
| Sun-OR Halogen | 0.6606 | 0.7813 | * |
|  | 0.9794 | 0.5702 | ** |

